# Phase 1 of the National Institutes of Health Preprint Pilot: Testing the viability of making preprints discoverable in PubMed Central and PubMed

**DOI:** 10.1101/2022.12.12.520156

**Authors:** Kathryn Funk, Teresa Zayas-Cabán, Jeffrey Beck

## Abstract

**Introduction:** The National Library of Medicine (NLM) launched a Pilot in June 2020 to: 1) explore the feasibility and utility of adding preprints to PubMed Central (PMC) and making them discoverable in PubMed, and 2) to support accelerated discoverability of National Institutes of Health (NIH)-supported research without compromising user trust in NLM’s widely used literature services.

**Methods:** The first phase of the Pilot focused on archiving preprints reporting NIH-supported SARS-CoV-2 virus and COVID-19 research. To launch Phase 1, NLM identified eligible preprint servers and developed processes for identifying NIH-supported preprints within scope in these servers. Processes were also developed for the ingest and conversion of preprints in PMC and to send corresponding records to PubMed. User interfaces were modified for display of preprint records. NLM collected data on the preprints ingested and discovery of preprint records in PMC and PubMed and engaged users through focus groups and a survey to obtain direct feedback on the Pilot and perceptions of preprints.

**Results:** Between June 2020 and June 2022, NLM added more than 3,300 preprint records to PMC (viewed 4 million times) and PubMed (viewed 3 million times) Nearly one-quarter of preprints in the Pilot were not associated with a peer-reviewed published journal article. User feedback revealed that the inclusion of preprints did not have a notable impact on trust in PMC or PubMed.

**Discussion:** NIH-supported preprints can be identified and added to PMC and PubMed without disrupting existing operations processes. Additionally, inclusion of preprints in PMC and PubMed accelerates discovery of NIH research without reducing trust in NLM literature services. Phase 1 of the Pilot provided a useful testbed for studying NIH investigator preprint posting practices, as well as knowledge gaps among user groups, during the COVID-19 public health emergency, an unusual time with heightened interest in immediate access to research results.

## Introduction

Scholarly communication, which encompasses the publication, dissemination, and discovery of research results [1], is a critical component of the biomedical research enterprise. As the largest public funder of biomedical research in the world [2], the National Institutes of Health (NIH) is committed to ensuring that the publications resulting from the research it funds are publicly accessible, widely disseminated, and broadly discoverable. This commitment is epitomized by the NIH Public Access Policy, established in 2008, which requires deposit of final, peer-reviewed manuscripts reporting NIH-supported research to be made publicly available in PubMed Central (PMC), the National Library of Medicine’s (NLM) digital archive for journals and articles, no later than 12 months after journal publication. In the ensuing 14 years, 1.4 million peer-reviewed articles with NIH support have been made available to the public in PMC under this policy, and discoverable in PubMed. More broadly and consistent with its mission, NLM supports public access to research outputs to accelerate scientific discovery and advance the health of individuals and our communities [3].

Since the establishment of the NIH Public Access Policy, scholarly communication has evolved as the number of journals and articles published annually has grown and new models of publishing have emerged. This growth has been accompanied by the emergence of new business models, the rise of open access publishing, increased attention to licensing terms and data sharing, and the increased use of preprints. The emergence of preprint servers and increased use of preprints in recent years has been described as, “Perhaps the biggest change in scholarly infrastructure” particularly, “in areas such as biology and chemistry where there had hitherto been little appetite for their take up” [4]. Though preprint posting in the biomedical and life sciences began to increase with the launch of bioRxiv in 2013, overall publication rates remained low compared to the journal literature [5].

Over the last few years, NIH has explored the role of preprints, which NIH defines as, “a complete and public draft of a scientific document … typically, unreviewed manuscripts written in the style of a peer-reviewed journal article,” [6] in sharing results of federally funded research [7]. A 2016 NIH request for information noted that “[p]reprints give their authors a fast way to disseminate their work, establish priority of their discoveries, and obtain feedback. Early-career scientists can also use preprints as evidence of independence and productivity.” Subsequently, in 2017 NIH began encouraging investigators to use preprints and other interim research products to speed the dissemination and enhance the rigor of their work [8]. However, preprints were considered out of scope for PMC and PubMed at the time because they were documents made public prior to peer review.

Preprints rose in prominence as a channel for rapid dissemination of biomedical research results during the COVID-19 pandemic [9]. Recognizing the benefits provided by accelerated discovery of preprints in these circumstances, on June 9, 2020, NLM launched the NIH Preprint Pilot (Pilot) to test the feasibility and utility of making preprints resulting from NIH-funded research available via PMC and discoverable in PubMed [10], consistent with NLM strategic efforts to “stimulate new forms of scientific communication and become the library of the future” and to “anticipate developments such as preprints” in scholarly communications [11].

This article describes the Pilot’s objectives, scope and approach, and summarizes findings to date.

## Objectives

The NIH Preprint Pilot was undertaken to inform NLM’s understanding of the role of preprints in scholarly communication and how they may fit into NLM literature services. NLM had two primary objectives in launching Phase 1 of the NIH Preprint Pilot:

1. To explore the feasibility and utility of identifying and archiving NIH-supported preprints in PMC with an associated citation in PubMed; and
2. To support accelerated discoverability of NIH-supported research results without compromising user trust in NLM’s widely used literature services.

## Methods

To launch the Pilot, we established its scope, identified eligible preprint servers, and developed processes for identifying and ingesting NIH-supported preprints. We leveraged PMC infrastructure to support the full-text archiving and indexing of all openly licensed preprints that were identified as within scope and to create metadata and abstract records for those preprints that were posted under more restrictive license terms. We also modified the PMC and PubMed user interfaces to enable users to differentiate between preprints and published articles on search results and article records.

Over the two-year Pilot, we collected data on retrievals of NIH-supported preprints and monitored changes in publication status of preprints. We also engaged users through focus groups and a survey to understand public perception of preprints and obtain direct feedback on their inclusion in PMC and PubMed.

### Scoping

NLM defined the scope of the Pilot as limited to preprints resulting from research conducted or funded by NIH (i.e., “NIH supported”). NLM considered that NIH procedures for selecting and monitoring research [12] would provide an important element of trust and help ensure quality as preprints make research results public prior to peer review.

To further narrow the scope of the Pilot, NLM focused Phase 1 on preprints reporting NIH-supported research relating to the SARS-CoV-2 virus and COVID-19. This limited the number of preprints included in Phase 1 and targeted a research area for which there was considerable interest in accelerated access to research results by a broad range of users, including researchers, clinicians, public health officials, and the general public. Although an atypical situation given the urgency of information access about a novel disease to inform immediate action, SARS-CoV-2 and COVID-19 research presented an active testbed for the Pilot.

### Selecting preprint servers

To identify preprint servers for inclusion in the Pilot, we applied three general criteria:

1. Public practices largely aligned with NIH guidance on preprint server selection [13] and emerging community practice [14], including:

○ policies regarding plagiarism, competing interests, and misconduct and other hallmarks of reputable scholarly publishing are rigorous and transparent;
○ records of changes are maintained, and users have clear ways to cite different versions;
○ maintaining links to the peer-reviewed journal version, if available;
○ publicly posted screening process; and
○ robust archiving strategy that ensures long-term preservation and access;
2. Likely to contain NIH-funded research; and
3. Indexed in the NIH Office of Portfolio Analysis iSearch COVID-19 Portfolio [15] at the time of the Pilot launch.

### Technical implementation

Technical implementation of the Pilot involved leveraging the existing PMC infrastructure for the ingest and archiving of articles and developing new processes for preprint identification and conversion, in addition to modifications to the PMC and PubMed user interfaces.

### Preprint identification

To identify preprints reporting NIH-supported SARS-CoV-2 virus or COVID-19 research, NLM established text mining processes to locate text strings that could be matched to NIH grants or contracts. A web interface was developed to support staff review and confirm accuracy of suggested text mining results.

To determine the relevance of research reported on SARS-CoV-2 virus or COVID-19, NLM relied on the NIH Office of Portfolio Analysis iSearch COVID-19 Portfolio tool. NLM also used this tool to identify preprints with NIH-affiliated authors (i.e., intramural researchers and staff).

Extramural and intramural preprint identification processes were conducted weekly.

### Ingest and conversion processes

Each week, following preprint identification, NLM staff upload a list of the permanent identifiers (mostly digital object identifiers or DOIs) for those preprints identified as reporting NIH-supported SARS-CoV-2 virus or COVID-19 research to a PMC tool developed for implementation of the Pilot. This triggers an initial ingest process that extracts title, author, and abstract metadata for those DOIs on the list into PMC. A PMC identifier (PMCID) is then assigned and a corresponding title and abstract record is loaded to PubMed.

NLM then converts the full text of those preprints made available under a Creative Commons license to archival XML for inclusion in PMC. All full-text content in PMC is stored in the most recent American National Standards Institute (ANSI) and National Information Standards Organization (NISO) Journal Archiving and Interchange Tag Suite (JATS) XML format, which is currently ANSI/NISO Z39.96-2021 JATS [16].

Those preprints identified as in scope for the Pilot but made available under other more restrictive license terms were included as metadata- and abstract-only records in PMC with links to the preprint server full text.

### Preprint record maintenance

Indexing and archiving preprints requires active record maintenance in PMC and PubMed. Scripts developed by NLM staff are run weekly to identify and ingest new versions of preprints in bioRxiv and medRxiv. All versions of a preprint share the same PMCID. PMC displays the most recent version of the preprint available; previous versions remain accessible through the “Other versions” link in PMC.

To connect users to the peer-reviewed journal version when available, NLM staff conduct additional automated checks across the following resources to identify peer-reviewed journal versions of preprints: bioRxiv API [17]; Crossref API [18]; Europe PMC RESTful API [19]; and PubMed Citation Matcher, an NLM-developed resource that compares the title, author lists, and abstracts of preprints with PubMed records.

We also established processes to check weekly for withdrawn preprints and subsequently ingest the withdrawal notice. In such cases, the title of the preprint in PMC and PubMed is updated to indicate the withdrawn status. NLM staff also run daily checks for retractions of journal articles, including those that have a corresponding preprint record in PMC and PubMed.

### Preprint record display

To conform with recommended community practice regarding preprints [20] and ensure a transparent scientific record, user interface modifications were made in PMC and PubMed to clearly identify preprint records as such, and provide links to preprint servers and, when available, associated peer-reviewed journal versions.

A prominent green information panel alerting the user that the record being viewed is a preprint was added to all preprint records in PMC and PubMed to distinguish them from journal article records. The text in this panel notes that the article has not been peer reviewed and includes a link to more information about the “NIH Preprint Pilot.” To communicate the “NIH-supported” scope of the pilot, an NIH-branded preprint banner was also added to records in PMC (Figure 1).

**Figure 1.**
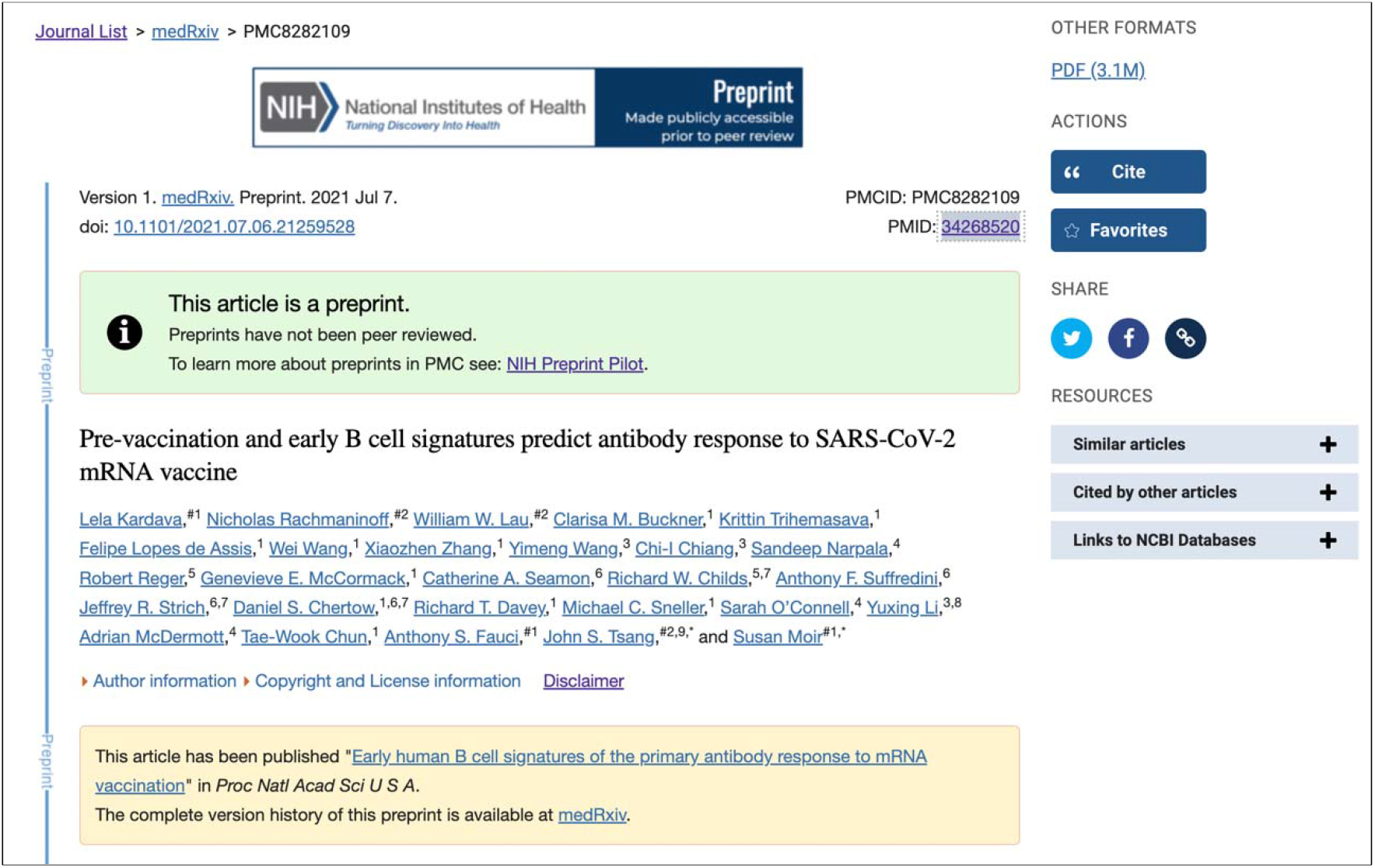
Screenshot of a preprint record display in PMC (PMC8282109). This example record includes green information panel identifying the record as a preprint that has not been peer reviewed, the preprint indicator in the citation, and the yellow related content information panel that points to the associated peer-reviewed journal version and preprint server.

A “Preprint” indicator was also added to the displayed citation metadata and “Cite” tool in PMC and PubMed to foster transparency as well as accurate citation (Figure 2).

**Figure 2.**
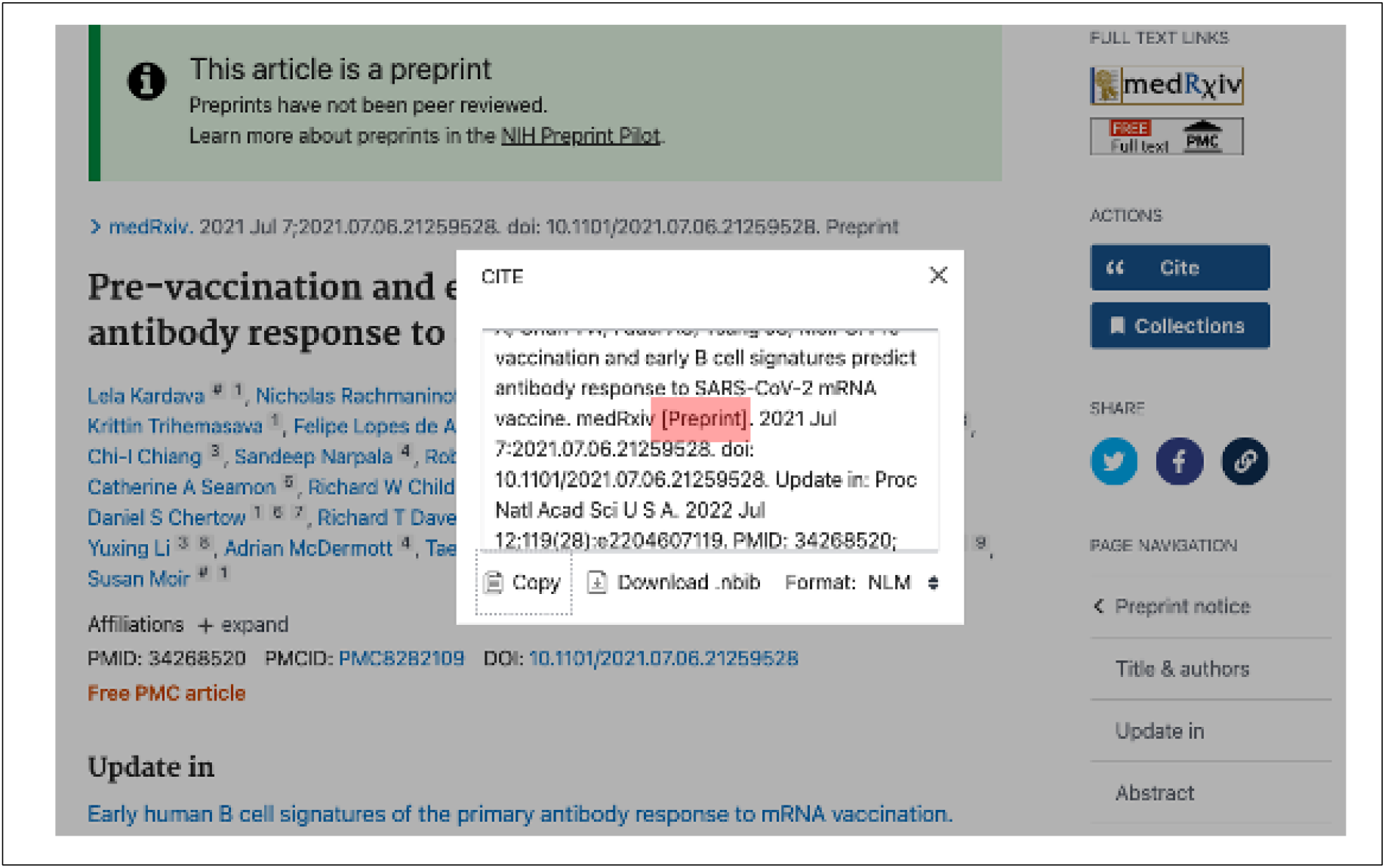
Screenshot of a preprint record display in PubMed (PMID: 34268520). This example record includes green information panel identifying the record as a preprint that has not been peer reviewed, the preprint indicator in the citation metadata, and Cite tool pop-up window with the “[Preprint]” indicator.

Additionally, the yellow information panel in PMC that displays prior to the abstract and includes related content links was expanded to include a pointer to the preprint on the source preprint server website and a link from the preprint record to an associated peer-reviewed journal version, when available (Figure 1). Users may also access and view the preprint record directly from the source preprint server by clicking on the server link in this panel, the hyperlinked DOI in PMC and PubMed, or the server-branded “LinkOut” button in PubMed (Figure 2).

We added similar “Preprint” citation indicators to preprint records in the search results of PMC and PubMed. Preprints that were linked to published journal articles were labeled as “Updated” in PubMed (Figure 3). In PMC, a “Published in” link was added to the search results display to take the user directly to the peer-reviewed journal version, if available (Figure 4).

**Figure 3.**
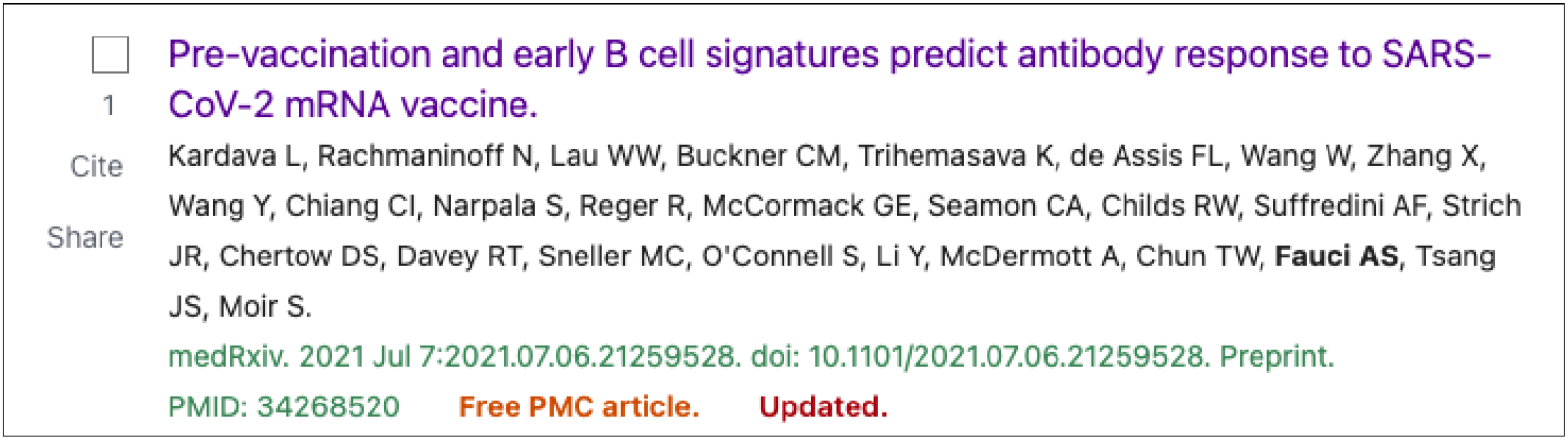
Screenshot of a preprint record search result in PubMed. This example includes the preprint indicator following the DOI as well as the Updated identifier, indicating that a peer-reviewed journal version is available.

**Figure 4.**
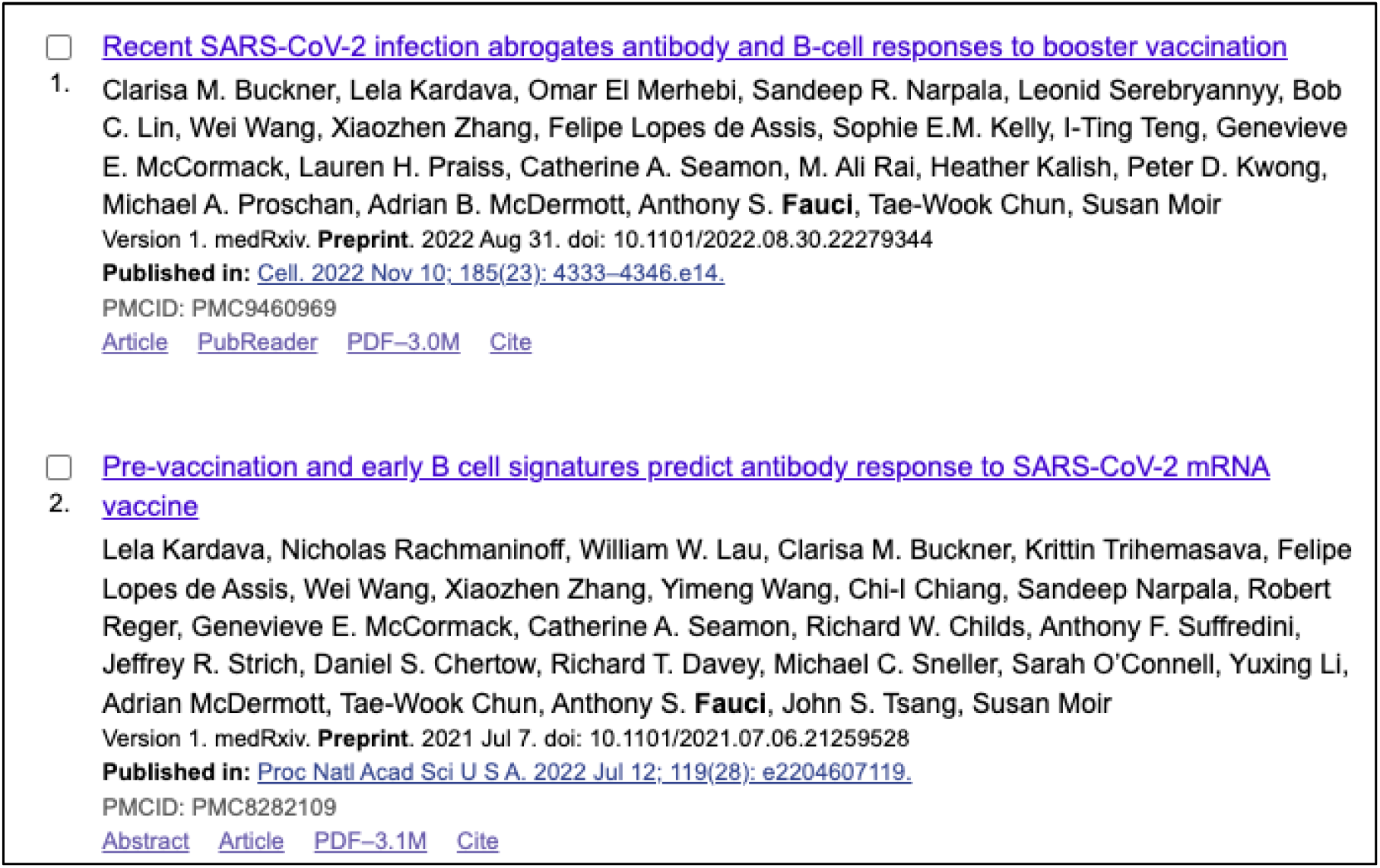
Screenshot of search results that include preprint records in PMC. These examples include the “Preprint” indicator following the preprint server name and display of the “Published in” link for any associated peer-reviewed journal version.

Finally, to enable easy identification of preprint records in PMC and PubMed in search processes, NLM created search filters. In PMC, users can apply the preprint[filter] to any search. In PubMed, users can search by publication type (preprint[pt]) or retrieve preprint records via E-utilities, using the publication type “Preprint”. These search filters also allow users to exclude preprint records from search results by using the Boolean “NOT” in either database, e.g., “covid 19 NOT preprint[filter]” in PMC and “covid 19 NOT preprint[pt]” in PubMed.

### Preprint use and practice monitoring

We used Google Analytics and internal web logs for preprint records to monitor preprint record use and engagement consistent with NLM Web Privacy and Security Policy [21]. Quarterly summary reports were published on the NIH Preprint Pilot webpage under the Related Links section [22] for public view. Throughout the Pilot, NLM monitored how preprint posting accelerated dissemination of and access to NIH-funded research via PMC and PubMed. NLM also monitored where NIH-supported SARS-CoV-2 virus and COVID-19 research was published as a preprint and under what license terms.

Recognizing that PMC as a full-text archive and PubMed as a citation and abstract database have unique roles to play in discovery, we implemented different methods for measuring how preprints were discovered in these databases. For each database, we extracted data from October 2021, a month that generally reflects use during the academic year. For PMC we compared usage of openly licensed preprints that include full text with preprint records that were citation and abstract-only in PMC. For PubMed, we examined the frequency that preprint records were returned in search results and viewed.

### User feedback

We also took steps to inform our understanding of the impact of the Pilot on public trust in NLM literature resources. Prior to launch, NLM established a preprint-specific email alias expressly for pilot feedback. Additionally, in summer of 2021, after the Pilot had been taking place for just over a year, NLM conducted focus groups and administered a survey to understand user perceptions on preprints and their inclusion in PMC and PubMed.

NLM conducted four online focus groups, with eight to nine participants per group. Represented were key user groups of NLM literature resources (biomedical researchers, clinicians, and research librarians) as well as healthcare journalists, a group that often acts as an intermediary between the research results that are published or made publicly available and the public. A nationwide consumer research company recruited the clinicians and the researchers. The healthcare journalists and medical librarians were recruited through the Network of the National Library of Medicine and existing NLM relationships. Participants with a mix of professional experience and familiarity with preprints were selected for each group to participate in a 2-hour discussion, conducted via Zoom (for more detail see focus group guides in Supplemental File 1).

In addition, NLM administered an online feedback survey (OMB Control No: 0925-0648) in August and September 2021, which was made available in PMC and PubMed to users who accessed preprint records in these databases. Surveying PMC and PubMed users allowed us to collect data on a broader set of user groups than those engaged in the focus groups, including students and educators, in the specific context of preprints in NLM databases. Because of the low overall numbers of preprint records in PMC and PubMed in comparison to journal article records, to survey database users that view preprint records we set high sampling rates. A feedback prompt was made available to 30% to 40% of users that viewed a preprint record in either PMC or PubMed during the 2-month period.

Only users that indicated previous knowledge or awareness of preprints were asked more detailed questions about their perspectives on preprints. The complete set of survey questions are available in Supplemental File 2.

## Results

### Discovery of NIH research

Between June 9, 2020 and June 9, 2022, NLM made more than 3,300 (n=3,332) preprint records discoverable in PMC and PubMed (see Supplemental File 3 for complete list). This represents approximately 8% of all preprint records reporting on SARS-CoV-2 virus and COVID-19 research included in the NIH Office of Portfolio Analysis iSearch COVID-19 Portfolio tool during that period (iSearch does not limit its portfolio to NIH-supported preprints). Under 10% (303) of these preprint records included NIH author affiliation data in PMC and PubMed. The majority were supported by an NIH extramural award and identified through text mining processes. Over the course of the Pilot, preprints have been viewed 4 million times in PMC. Corresponding preprint records in PubMed for all preprints ingested into PMC have been viewed more than 3 million times.

NLM included the following preprint servers in the Phase 1 based on eligibility criteria: medRxiv, bioRxiv, Research Square, arXiv, ChemRxiv, and SSRN. Of the preprint records added to PMC and PubMed, the majority were posted to either medRxiv (47%) or bioRxiv (38%) (Figure 5).

**Figure 5.**
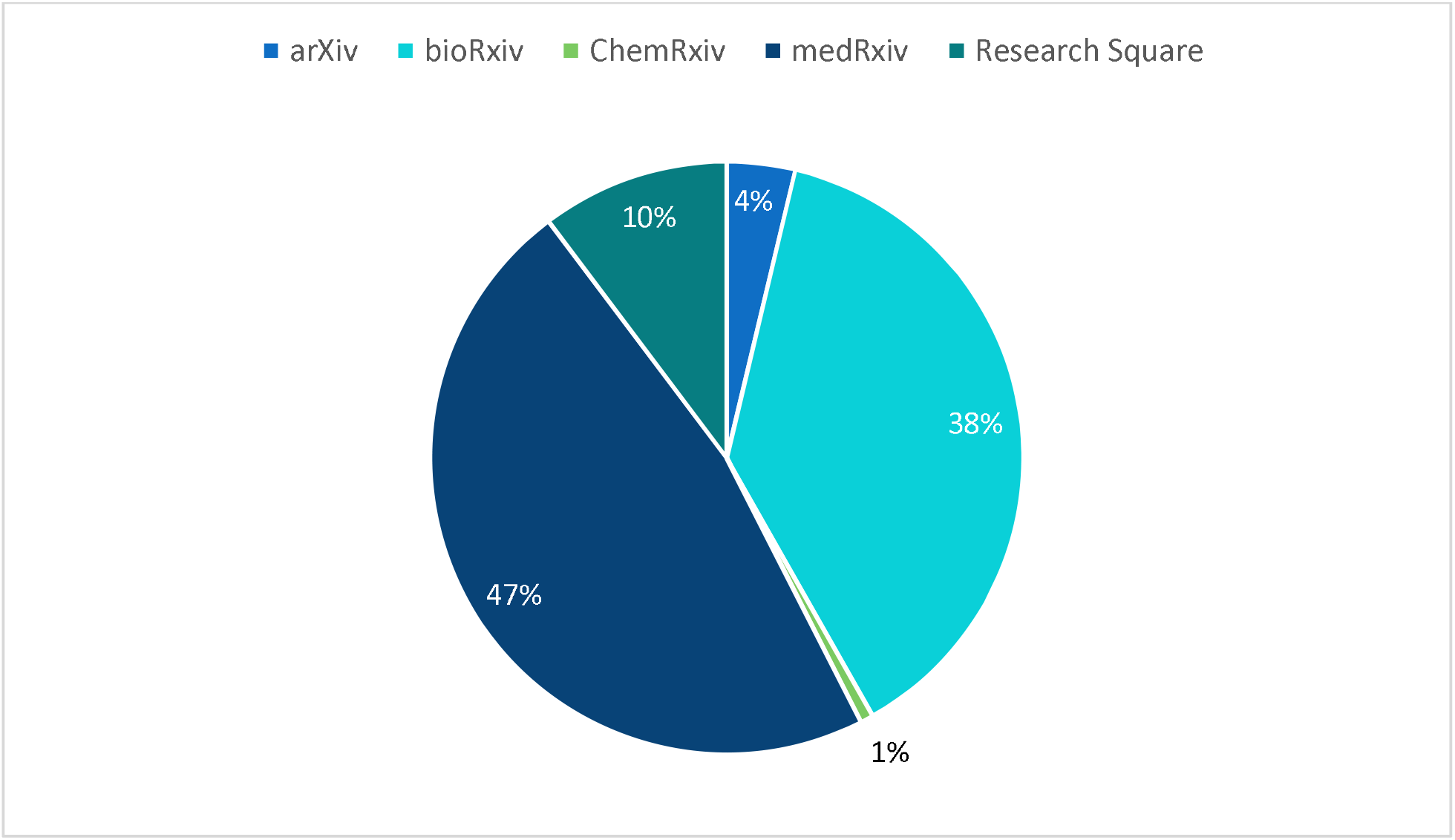
Breakdown of preprints by server during the first two years of the Pilot (see Supplemental File 3).

In 2021, the NIH Office of Portfolio Analysis expanded the scope of its iSearch COVID-19 Portfolio to include preprints posted to preprints.org and Qeios. Analysis completed by NLM did not find a sufficient volume of NIH-funded preprints in either of these servers to merit setting up new curation and ingest processes to include these preprint servers in Phase 1.

The volume of preprints identified as in scope for Phase 1 varied over time, peaking at 538 preprints in the first quarter of the pilot (June 9 – September 9, 2020; see Figure 6). Only one preprint was withdrawn by the authors or preprint server during Phase 1. To date, no preprints included in Phase 1 have been retracted following publication in a journal.

**Figure 6.**
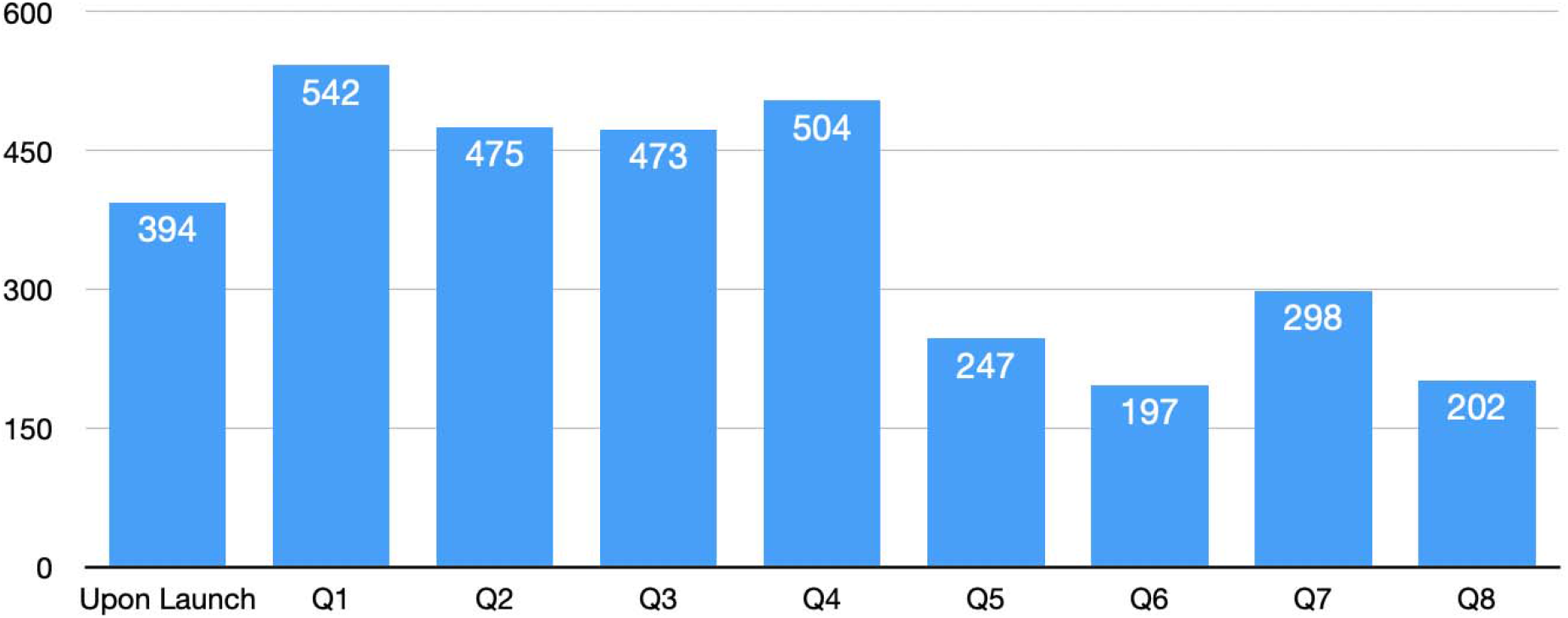
Number of preprints added to PMC upon launch (June 9, 2020) of the NIH Preprint Pilot and in each subsequent quarter of the pilot during the first 2 years (see Supplemental File 3).

Some preprint servers included in the Pilot (e.g., Research Square) require authors to apply a Creative Commons license. Others, such as bioRxiv and medRxiv allow authors to select from a “menu” of license options, ranging from traditional copyright restrictions to Creative Commons with attribution or CC0/public domain for U.S. government employees.

Since June 2020, there was quarterly growth in the number of NIH-supported authors selecting some type of Creative Commons license (see Figure 7). More commonly NIH-supported authors selected the more restrictive Creative Commons license options when available, limiting use to noncommercial reuse and no derivatives, or to make the work available under traditional copyright restrictions. This, in turn, limits what is archived in full-text XML in PMC.

**Figure 7.**
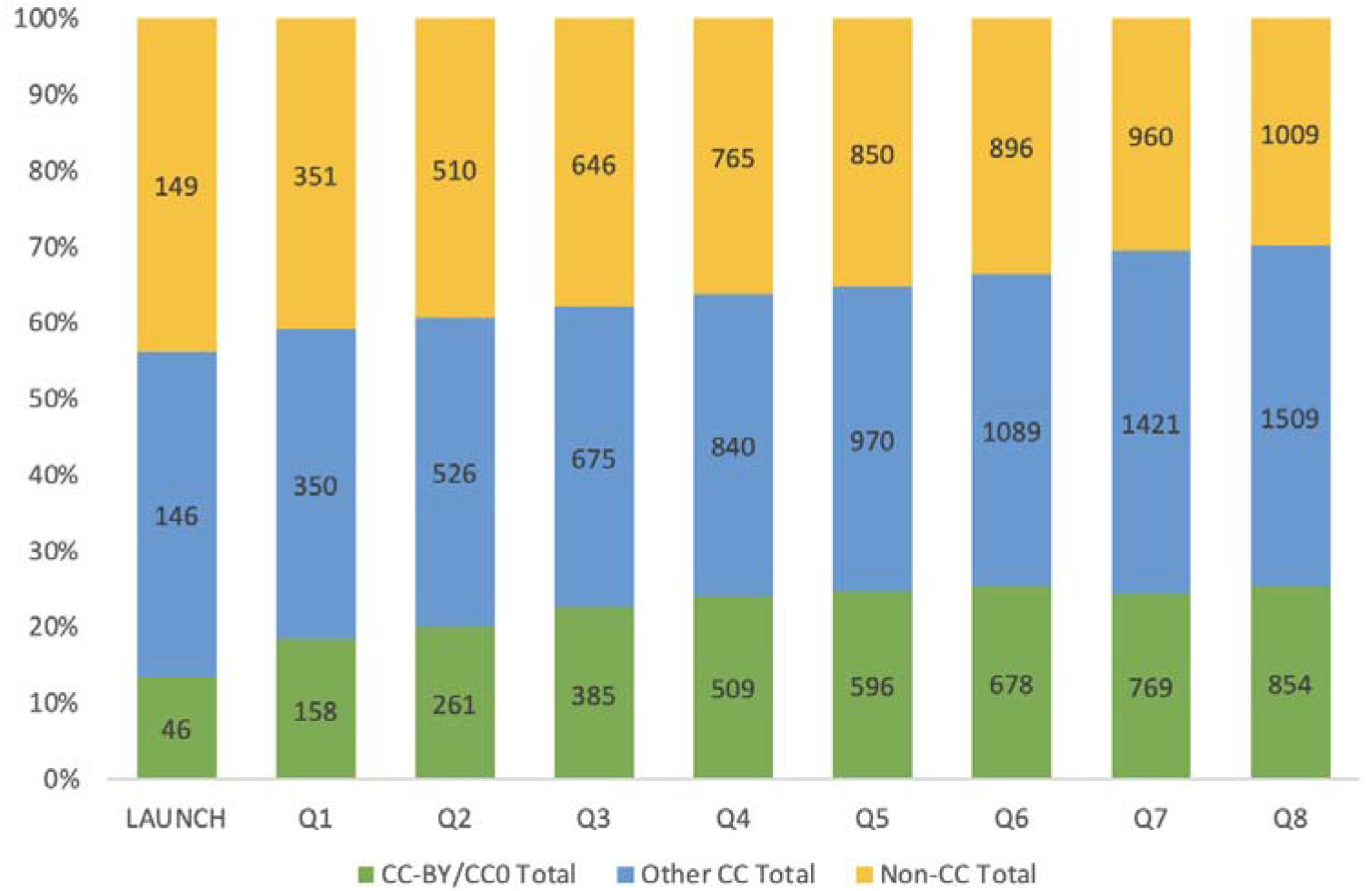
Bar graph showing number of preprints added to the Pilot at the time of launch and in each subsequent quarter of Phase 1, by license type.

PMC usage data from October 2021 for bioRxiv preprints added to PMC in 2021 (*n* = 482) were analyzed to inform our understanding of the role of full text availability in discoverability of preprints, as bioRxiv is one of the preprint servers that includes a mix of Creative Commons licensed preprints and preprints under traditional copyright. Therefore, the sample included a mix of openly licensed preprints with full text available and restricted licensed preprints that were available as citation and abstract records only. Preprints in the sample available under a Creative Commons license had on average been available in PMC for 190 days; preprints in the sample made available under more traditional copyright restrictions had been available in PMC for an average of 191 days. The data presented in Table 1 compares the Unique User IPs that accessed preprints in this sample during October 2021 and illustrates the overall higher rates of unique user engagement in PMC with preprints made available under a Creative Commons license.

**Table 1.** Analysis of generalized, aggregate data on unique user IP engagement with preprints in PMC during October 2021 that have full text available (yes – Creative Commons License) vs. those that are metadata-abstract records only (no – Creative Commons License).

|  | Minimum | 1<sup>st</sup> Quarter | Median | Mean | 3<sup>rd</sup> Quarter | Maximum |
| --- | --- | --- | --- | --- | --- | --- |
| No<br>Creative<br>Commons<br>License (*n*<br>= 171) | 1.00 | 8.00 | 12.00 | 16.14 | 20.00 | 95.00 |
| Has<br>Creative<br>Commons<br>License (*n*<br>= 311) | 2.00 | 14.00 | 25.00 | 41.43 | 42.50 | 367.00 |

Additionally, NLM found that 98.4% of all available preprint records in PubMed were viewed by users and that 99.4% of available preprint records were returned in search results during October 2021, reflecting the demand for research on the SARS-CoV-2 virus and COVID-19. The 17 records (0.6%) that were not returned were added to the database at the end of the timeframe analyzed, which is the likely reason for their absence from search result data. Of the 2,767 preprint records available in PubMed at the end of October 2021, there were only 20 that were returned in search results that were not viewed.

### Accelerated discovery

Approximately 72% of or 2,512 preprints added to PMC and PubMed through June 2022 had been linked to a peer-reviewed journal version by December 2022 (Figure 9). Analysis completed a year into the Pilot compared the preprint posting dates of nearly 800 preprints in the pilot at the time, to the publication date of a linked journal article found that on average 100 days elapsed between preprint posting and journal publication. The maximum time elapsed between preprint posting and publication in this sample was 365 days. Repeating this analysis on sample data from the second year of the Pilot in June 2022 found an increase from an average of 100 days to 162 days from preprint posting to journal publication.

**Figure 9.**
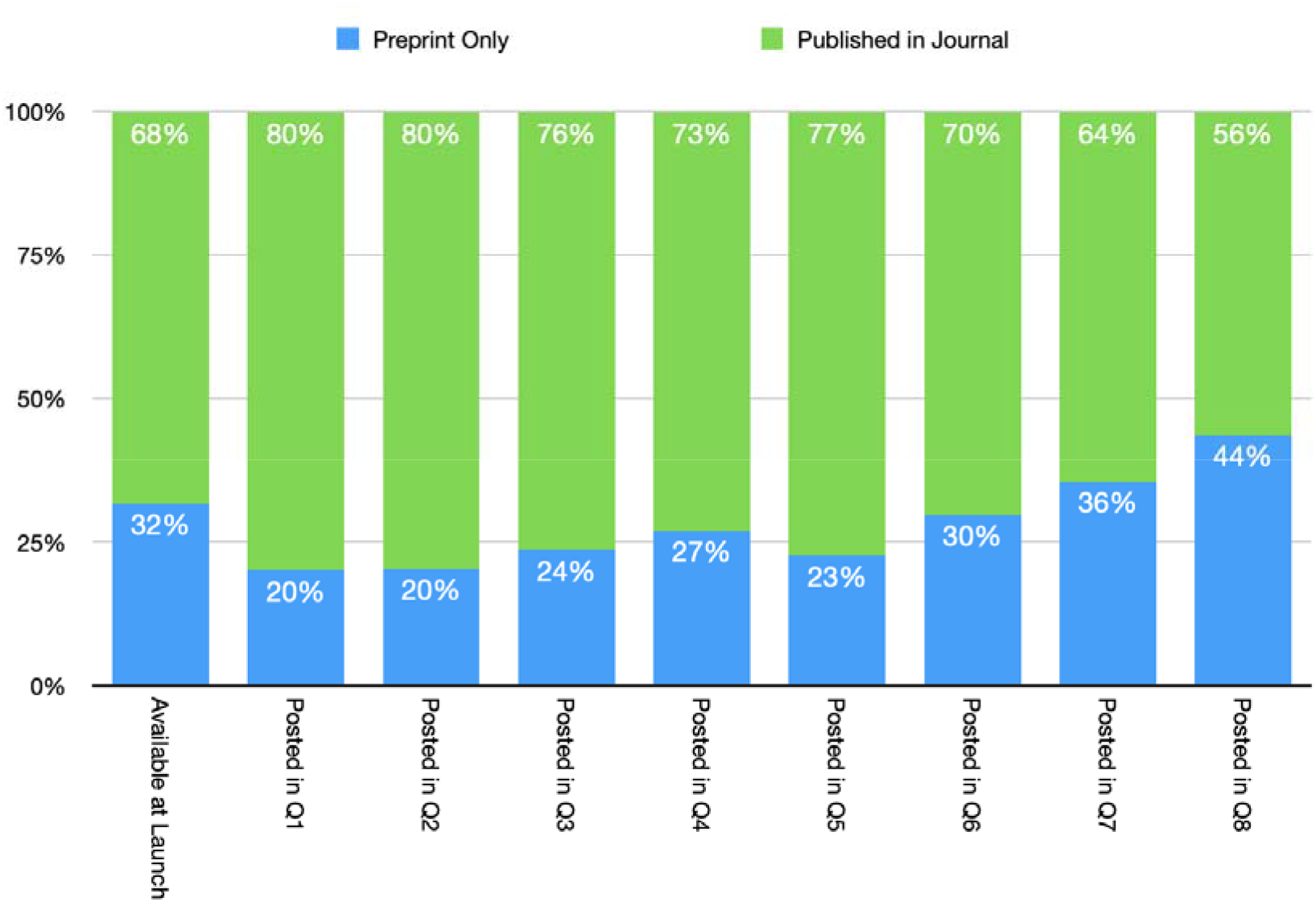
Quarterly breakdown of preprint status as of December 2022 based on date preprint was posted.

Of the journal articles linked to preprint records added to PMC during the first 2 years of the pilot, approximately 90% or 2,292 of those published articles were publicly accessible in PMC. This high proportion of publicly available journal articles in PMC is primarily due to the open availability upon publication of journal articles reporting relevant research and deposited in PMC as part of the PMC COVID-19 Collection [23].

### Attitudes toward preprints in NLM literature services

#### Email feedback

During the first year of the Pilot, 50 individuals contacted the NLM preprint email address; an additional 10 individuals either contacted NLM staff directly via email or used another NLM email address to provide feedback on the Pilot (See Supplemental File 4). The most common type of feedback received by the NLM preprint email address were requests by authors to add a preprint to PMC and PubMed (*n* = 28). Nineteen individuals had general questions about Pilot implementation, ranging from scope to version management to assignment of PubMed and PMC identifiers.

Seven of the 60 individuals that contacted NLM via email shared concerns. These concerns focused on:

- The perception or possibility of low-quality content being added to PubMed;
- Concerns about public understanding of preprints; and
- The potential impact on the reputation of NLM literature services.

Two concerns were received about the content of individual preprints associated with extramural projects. In both cases, the concerns were shared with the NIH program officer for the project, and in both cases, no issues were found with the preprints. One other email noted the lack of communication about NLM plans prior to the Pilot.

Overall, feedback received via email indicated:

- Authors are supportive of preprint discovery in PubMed and PMC.
- Authors would like the peer-reviewed journal version to be prioritized in discovery once available.
- Authors occasionally needed clarification on the scope of the Pilot.
- Not all users want to see preprint records in their search results.
- Need for clear and early communication of NLM plans that affect PMC and PubMed.

#### Focus groups

Focus group discussions provided NLM with qualitative data on:

- how different PMC and PubMed user groups (researchers, clinicians, medical librarians, and healthcare journalists) assess the content of articles and preprints;
- how they seek out and/or use preprints;
- how the pandemic influenced their perception of preprints;
- how they learned about preprints and what they suggest for use going forward; and
- the role of NIH and/or NLM in the proliferation of preprints.

From these discussions, we learned that research articles are assessed similarly by user groups. In assessing articles, participants considered the journal, magazine, or publication it appeared in; the author who wrote it; and the publisher. Other considerations included whether the publication is indexed in PubMed.

When the topic shifted to preprints, four themes emerged across groups: confusion, curiosity, caution, and an interest in or desire to be collaborative. As preprints are still a relatively new type of scholarly output in the biomedical and life sciences, most participants acknowledged that prior to the invitation to participate in the focus groups, they had not given preprints much thought or attention. Through further discussion, we found that:

○ When familiar with the concept of peer review, defining a preprint as “a document that has not yet gone through peer review,” is clear and understood.
○ Participants across all groups wanted to know what safeguards were in place to ensure that preprints would not be confused with vetted, peer-reviewed articles.
○ Researcher participants were interested in the potential of preprints as a new way to share research results.
○ Medical librarians, researchers, and healthcare journalist participants expressed that preprints were valuable; in particular, to learn about research results related to emerging topics, and to garner early feedback to improve reporting. However, some clinicians expressed doubts about the value of preprints to them and their work.

#### Survey

NLM received 321 responses to the survey between August and September 2021. Survey respondents represented a cross-section of PMC and PubMed user groups, with one-third of respondents being researchers (Figure 10).

**Figure 10.**
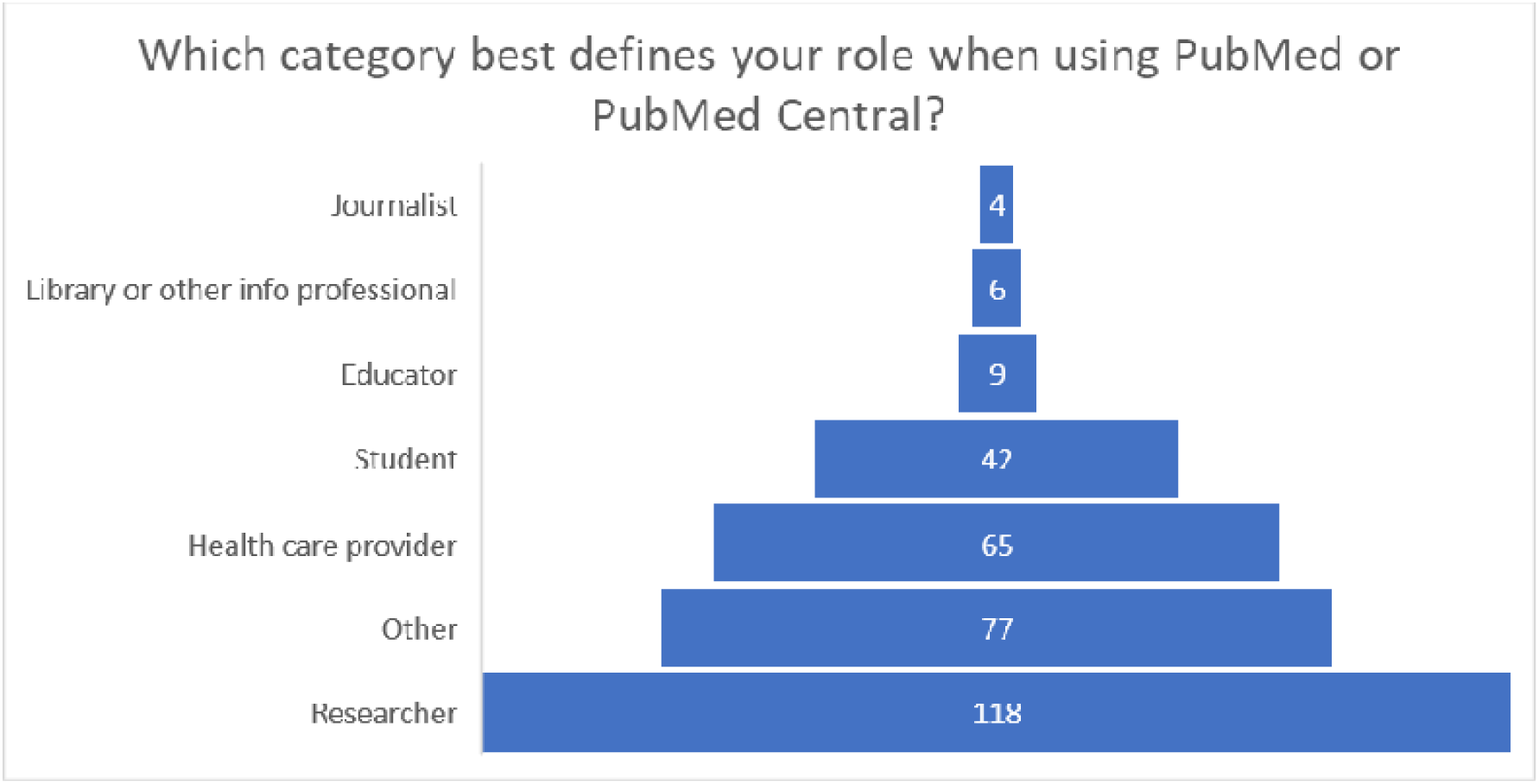
Number of respondents to the preprint survey by user group.

This survey collected quantitative, contextual data from those users engaging with preprint records in NLM databases to complement the qualitative data from the focus groups and inform our understanding of:

- Which user groups were accessing preprint records;
- The clarity of NLM’s presentation of preprints;
- Users’ awareness of preprints;
- Users’ confidence level in assessing scientific rigor of an article;
- The effectiveness of the preprint record display in communicating the type of content; and
- Users’ attitudes around the inclusion of preprints in PMC and PubMed.

NLM was also interested in how the availability of COVID-related preprints in PMC and PubMed affected public trust of NLM and its literature services and identify user knowledge and skills gaps related to preprints.

Sixty-two percent of survey respondents (201) reported having previously heard of preprints, although there was variability across user types (see also Figure 11):

- Seventy-five percent of researcher respondents reported they had heard of preprints.
- Educator (67%) and student (60%) respondents reported being somewhat familiar with preprints.
- Healthcare provider (57%) respondents reported that they were less familiar with preprints than researchers, educators, and students.
- Approximately one-half of other user (52%) respondents reported that they had heard of preprints.

**Figure 11.**
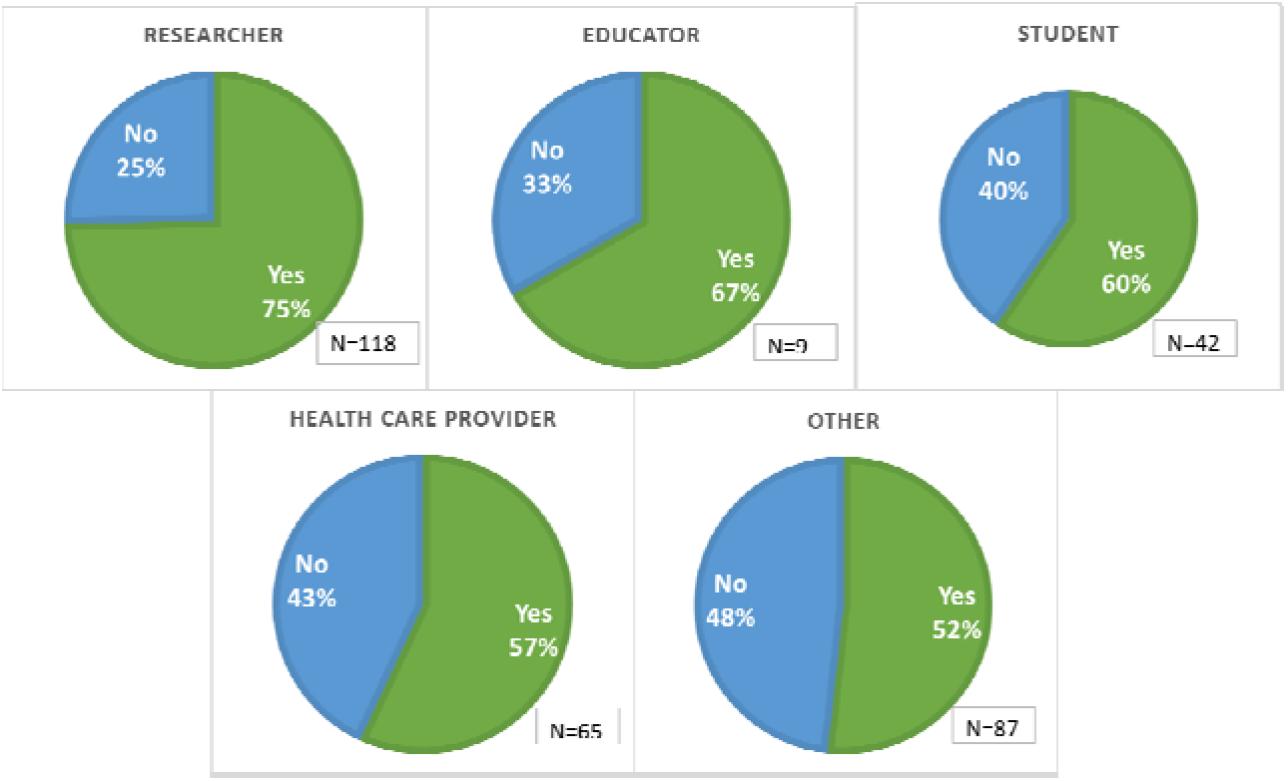
Survey respondents and their responses to whether they had previously heard of preprints in a survey of PubMed and PMC users in July and August 2021.

Survey responses indicated that users were generally able to distinguish that they were viewing a preprint record in PMC and PubMed; Seventy percent (177) indicated that it was “Clear,” noting differences across user types as shown in Figure 12.

**Figure 12.**
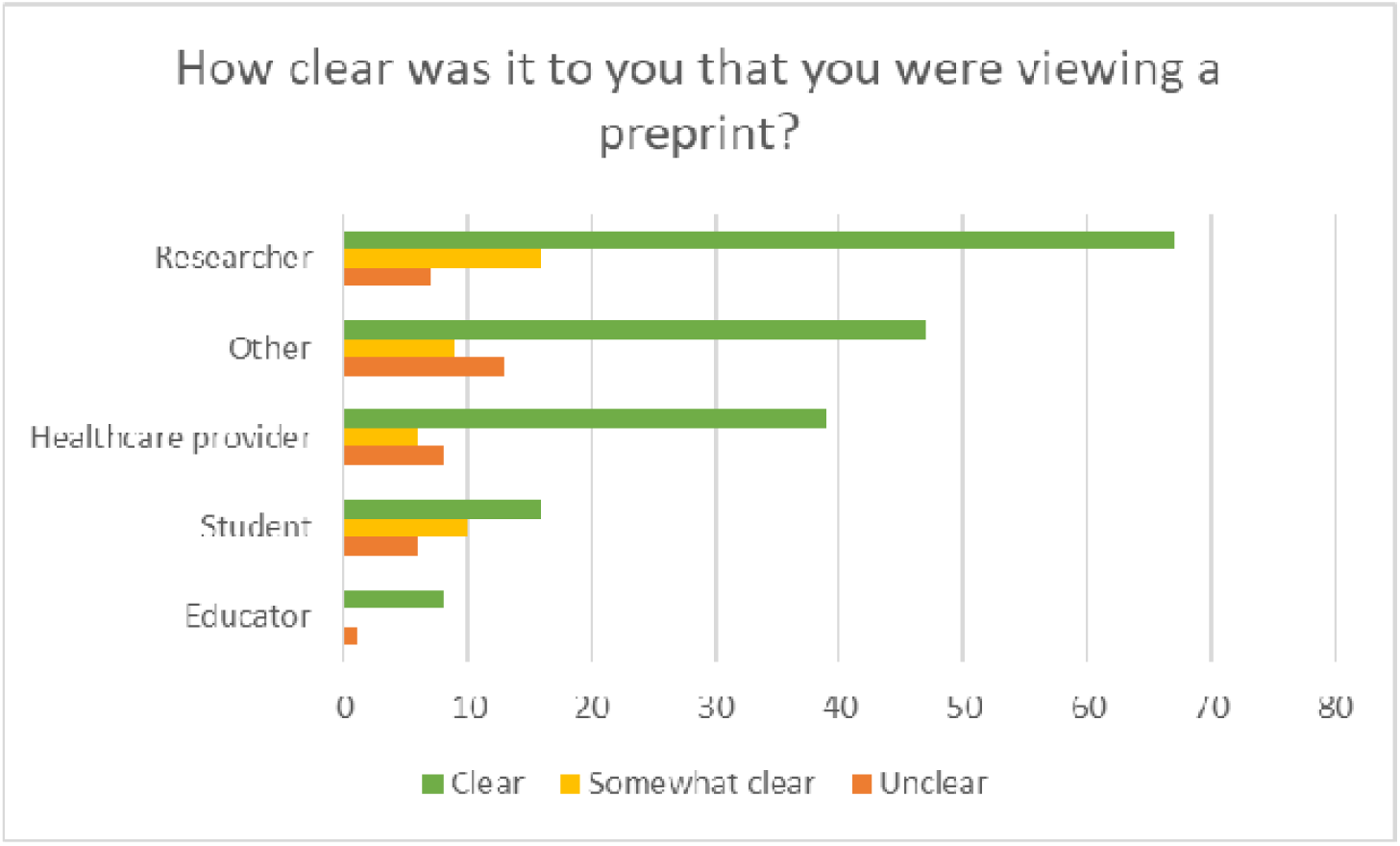
Users’ responses on whether it was clear to users that they were viewing a preprint, by audience type.

On average, survey respondents indicated that preprints are very important, especially for the scientific community at large, and even more so for emerging topics like the SARS-CoV-2 virus and COVID-19 (see Figure 13). Ninety-six percent of respondents felt that the scientific community’s ability to discover and access preprints was at least moderately important, and 92% of respondents felt that it was at least moderately important to discover and access preprints via PMC or PubMed.

**Figure 13.**
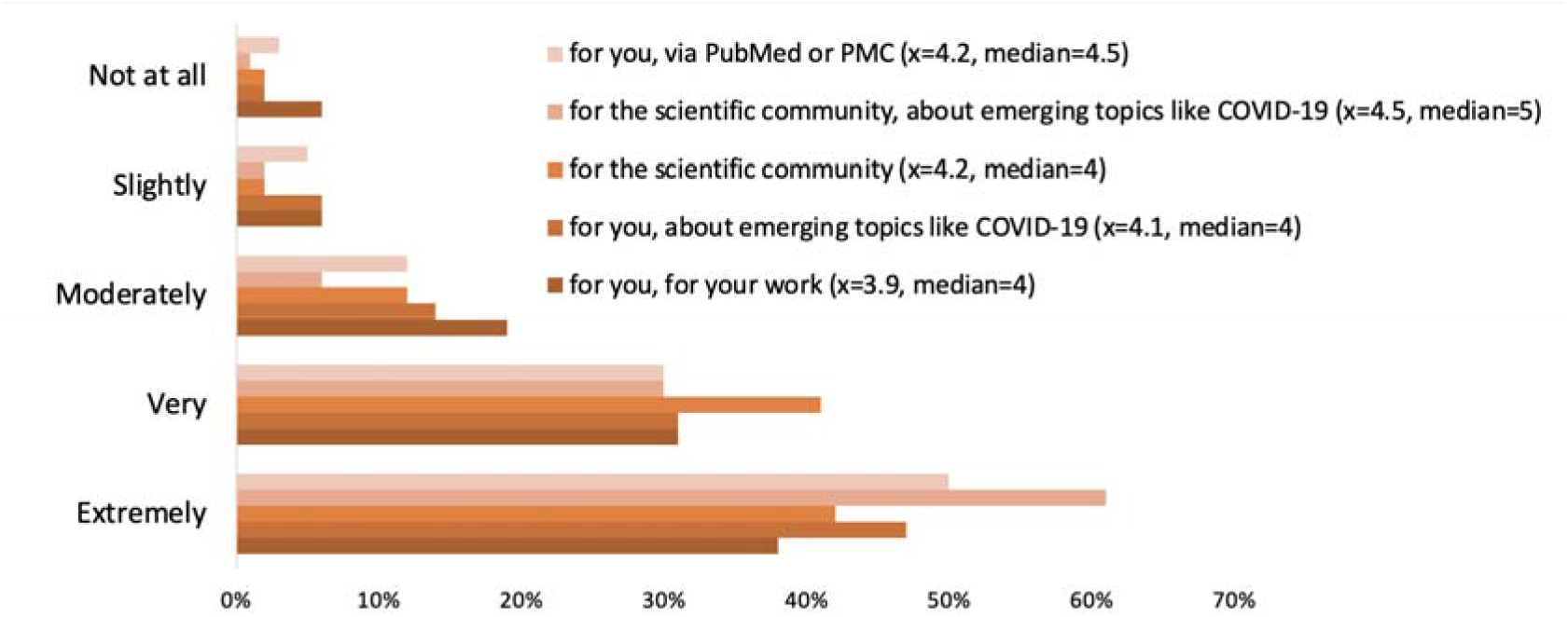
Survey responses on the importance of preprint discovery (i.e., “How important is it to be able to discover and access preprints?”)

Respondents reported that preprints make research results available more quickly, provide more exposure to research findings, and have the potential of improving the quality of the final product through wider review.

Fifty-seven percent (95) of survey respondents reported that having preprints in PubMed and PMC did not impact their trust in these databases, but for those for whom it did, it was more likely to increase than decrease their trust (see Figure 14).

**Figure 14.**
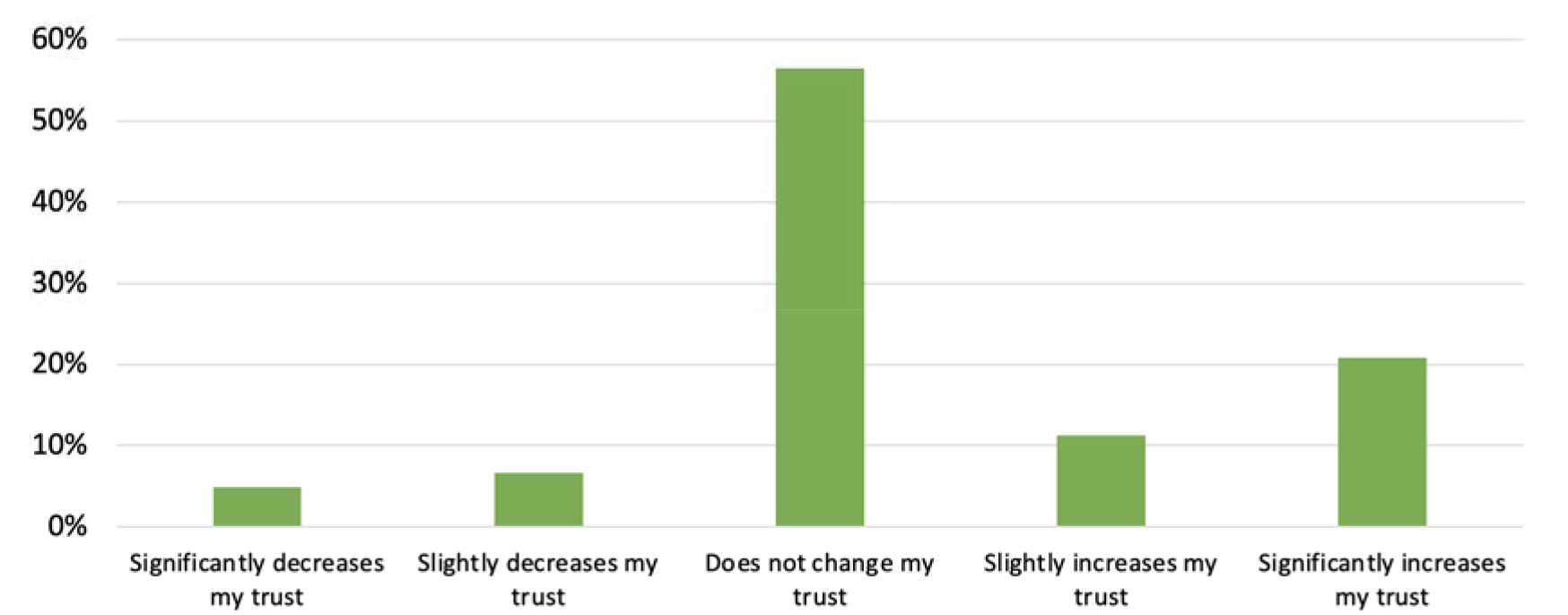
Responses to, “To what extent does having preprints in PubMed or PubMed Central increase or decrease your trust in the information found in PubMed or PubMed Central?”

For those whom it increased trust, respondent comments mentioned that their trust was gained by transparency. One respondent wrote, *“it shows willingness to present all ideas based on some scientific effort to be available for scrutiny by all.”* Of those respondents (12%/19) that noted availability of preprints decreased their trust in PubMed and PMC, one commented, *“I think of a library as a place where things are in final form.”*

See Supplemental File 5 for the complete response data.

## Discussion

During Phase 1 of the NIH Preprint Pilot, we confirmed the technical feasibility of leveraging existing NLM database infrastructure to ingest preprint records in PMC, and subsequently make them discoverable in PubMed.

Phase 1 allowed NLM to successfully test strategies for the identification of preprints that report NIH extramural and intramural research at a small scale. In particular, the work of the NIH Office of Portfolio Analysis proved to be key in preprint identification processes for intramural research as author affiliation metadata associated with preprints in machine-readable format was often not detailed enough to enable this type of identification otherwise.

Though several authors reached out to notify of us when in-scope preprints were missed in our identification processes, which were subsequently added to PMC and PubMed, there were no reports of preprints being inaccurately identified as NIH supported. Confirming that we were able to accurately identify those preprints within the scope of Phase 1 was a priority in implementation because we view the presence of NIH support as a key safeguard to the inclusion of papers made public prior to peer review in our databases. As noted by users, there are perceived risks regarding discoverability of the nonpeer-reviewed literature. The lack of controversy or validated concerns to date regarding preprints added under the Pilot demonstrate the value of keeping the scope consistent with NIH guidance and NLM collection guidelines.

Clear presentation and labeling of preprints as not peer reviewed was also prioritized in implementation. Though survey results indicate that preprint record labeling in PMC and PubMed was largely effective, there remain knowledge gaps across different user groups and some author emails indicate that more could be done to increase the clarity of the scope of the Pilot.

Since June 2020, much has been studied and written about the role of preprints in communicating COVID-19 research results. Otridge J et al. [24] found that “the incorporation of high-quality preprints into the CDC COVID-19 Science Update improve[d] this activity’s capacity to inform meaningful public health decision-making.” Similarly, during Phase 1, NLM found incorporating preprints within the larger corpus of curated scholarly literature made available in PMC and PubMed helped NLM contextualize the research reported in preprints, linking them to similar articles in PubMed, related data, and the larger record of citation in PMC.

Phase 1 of the NIH Preprint Pilot also supported accelerated discovery of NIH-supported research results. The analyses of PMC and PubMed usage data during Phase 1 informed our understanding of different paths users take to preprint discovery in NLM literature databases. These data also illustrated the value of indexing preprints in multiple resources that are integrated into different users’ literature search and discovery methods in different ways. As we saw, human-and machine-readable full text in PMC resulted in high rates of preprint discovery through third-party search engines. This was evidenced by higher overall preprint views in PMC than PubMed, more than two-thirds of which were from searches run outside NLM databases. Conversely, higher rates of PubMed users came to preprint records through a direct database search than those in PMC, signaling that preprints in PMC and PubMed may reach different users, depending on a user’s preferred search platform.

Additional data on PubMed user behavior showed that PubMed users were most likely to navigate from PubMed to the preprint server directly either via the DOI link or the LinkOut button to view the full text of a preprint, rather than to PMC. This again demonstrates that users may discover and engage with PMC and PubMed in different ways, with each playing a role in the wider information landscape.

Phase 1 results also highlight how even during a period of accelerated peer review and immediate open sharing of COVID-19-related literature, the indexing and archiving of preprints can speed the dissemination and discovery of NIH-supported research in PMC and PubMed. Specifically, we found that inclusion of preprints in PMC accelerated access to NIH research results in NLM literature databases by more than 100 days on average, a notable period of time during a public health emergency. In addition, their inclusion broadened access to NIH research, supporting discovery of NIH research results in our databases to nearly 1,000 articles that had not yet been published in a peer-reviewed journal or may not have been intended for formal journal publication.

A richer understanding of the characteristics of preprints in this latter “unpublished” subset, particularly a year or more after posting, and author motivations in preprint posting could contribute to a more complete picture of the role of preprints in offsetting publication bias and in communicating the results of federally funded research. How preprints may enable the sharing of nontraditional results (such as works in progress or negative, confirmatory, or contradictory results) is a topic that is largely unexplored in the current literature, though efforts to encourage the use of preprints for such purposes are emerging [25].

Though some focus group participants expressed general concerns about the quality of scientific literature made publicly accessible as a preprint prior to peer review, NLM did not find evidence to support these concerns for those preprints in scope for Phase 1. During this phase, no verified concerns about the quality of scientific reporting of any preprint added to PMC or PubMed were raised by users of these databases. This may be, in part, because NLM limited preprint collecting activities to those that report NIH support and, therefore, are subject to the NIH grant selection peer review process [26] or internal approval processes. NLM’s experience with Phase 1 is also consistent with results reported in a growing body of literature comparing the content of articles posted as preprints to the content of the same article following publication in a peer-reviewed journal [27–30]. For example, Nelson et al. found that “[o]verall, articles submitted to preprint servers by researchers, especially on COVID-19, are largely complete versions of similar quality to published papers and can be expected to change little during peer review” [31].

Phase 1 has further informed our understanding of NIH researcher preprint practices in the context of broader scholarly communication activities during the COVID-19 pandemic. More than 13,000 journal article records with NIH support reporting COVID-19-related research were added to PubMed during this period; about 10% of which were linked to a preprint record. A 2021 report found that only 5% of peer-reviewed articles reporting COVID-10 research had a corresponding preprint posted prior to journal publication peer-reviewed journal [32]. To what degree the Pilot may have impacted rates of preprint posting among NIH investigators is unknown. Further, though there have been steady increases in the number of openly licensed preprints added to PMC each quarter, we are unable to identify the impact the Pilot may have played in raising awareness of NIH recommendations on licensing nor the impact that culture of openness around COVID-19 research may have impacted author decisions around what license to choose when presented with options.

Our efforts to enable accelerated discovery of SARS-CoV-2 virus and COVID-19 research in PMC and PubMed, both as preprints and published articles, have increased our awareness of the challenges that come with archiving and presenting an archival scientific record that may include multiple versions of a paper and the importance of transparency as to the status and source of all records in our databases. As a first step, we developed and released the NIH Preprint Pilot Toolkit [33], an online resource for librarians to learn about the Pilot and preprints, as well as Preprints: Accelerating Research [34], an on-demand training, though more direct materials available directly from preprint records in PMC and PubMed may prove to be beneficial to certain user groups. With this in mind, we continue to review the presentation of preprints in PMC and PubMed to identify new strategies for communicating the status of preprints, facilitate connections between them and journal articles, and clearly convey the NIH-funded scope of our collection efforts.

Finally, while the focus groups and survey played a key role in identifying in knowledge gaps around preprints as well as informing NLM’s understanding of user perceptions on accelerated discoverability of NIH research result in PMC and PubMed, there are limitations on those findings. Focus groups only included a limited number of participants by user group. Survey respondents were interacting with a preprint record when the survey triggered, which may suggest either a willingness to engage with preprints, generally, as well as a likely interest in SARS-CoV-2 virus or COVID-19 research, or perhaps a lack of awareness of preprint status. As such focus group and survey findings may not be an accurate representation of user perspectives across research disciplines and specialties and thus may not be generalizable to all users or audiences, nor to all NIH-supported preprints.

## Conclusions

The NIH Preprint Pilot has confirmed the technical feasibility of including preprints in PMC and PubMed. Further, NLM has found that preprint records in PMC and PubMed provide an additional avenue for accelerated discovery of NIH-supported research during the ongoing public health emergency prior to journal publication. In addition, the Pilot did not have strong impact on customers’ trust of NLM and its literature services. In cases where users did report it having an impact, they indicated it was more likely to increase their trust due to the greater transparency.

Through the Pilot, NLM has accelerated and expanded broad discovery of publicly funded research results, helped maximized the impact of NIH funding, accelerated the point at which this research would otherwise be discoverable and publicly accessible in PMC and PubMed, and supported the NIH response to the public health emergency. Given the success of Phase 1, NLM launched Phase 2 in January 2023, expanding the scope of preprints eligible for inclusion in PMC and PubMed to any preprint reporting on NIH-funded research posted to those servers from Phase 1 that contained the highest volume of preprints reporting on NIH-supported research. Phase 2 is expected to last for at least a year to further inform NLM’s understanding of the role of preprints in disseminating NIH research [35].

Though peer review remains integral to scholarly communication, preprints are positioned to play an expanding role, notably in the distribution and discoverability of research, as awareness of preprints continues to grow, new publishing models incorporate preprints, and the potential of preprints to facilitate greater sharing of research results faster is realized. Clear guidance accompanied by active engagement with investigators could help build on lessons that have been learned during the NIH Preprint Pilot and throughout the public health emergency about accelerated open access to research results.

As the world’s largest biomedical library, NLM is uniquely positioned to provide discovery tools and to engage with the wider medical and public library communities to raise awareness of preprints, encourage education and training, and continually improve the presentation and integration of preprints into the wider scholarly record. With such efforts, NLM aims to enable transparency and rebuild public trust in science.

## Version History

This document was first made public at the preprint server bioRxiv on December 13, 2022 (<u>10.1101/2022.12.12.520156</u>). The current version has been updated to include Table 1, cleaned data in figures 5 and 6, and refined reporting of email feedback data during the first year of the pilot, adding supplemental data files for further reading. We also expanded discussion on the quality of scientific reporting in preprints and updated the status of the Pilot as of January 2024.

## Data Availability Statement

Underlying data on email feedback and survey results are available as supplementary information.

Additionally, details of the PubMed query run on 11 August 2022 to identify non-preprint COVID-19 related literature are as follows: (((“covid 19”[All Fields] OR “covid 19”[MeSH Terms] OR “covid 19 vaccines”[All Fields] OR “covid 19 vaccines”[MeSH Terms] OR “covid 19 serotherapy”[All Fields] OR “covid 19 serotherapy”[Supplementary Concept] OR “covid 19 nucleic acid testing”[All Fields] OR “covid 19 nucleic acid testing”[MeSH Terms] OR “covid 19 serological testing”[All Fields] OR “covid 19 serological testing”[MeSH Terms] OR “covid 19 testing”[All Fields] OR “covid 19 testing”[MeSH Terms] OR “sars cov 2”[All Fields] OR “sars cov 2”[MeSH Terms] OR “severe acute respiratory syndrome coronavirus 2”[All Fields] OR “ncov”[All Fields] OR “2019 ncov”[All Fields] OR ((”coronavirus”[MeSH Terms] OR “coronavirus”[All Fields] OR “cov”[All Fields]) AND 2019/11/01:3000/12/31[Date - Publication])) AND “nih”[Grant Number]) NOT “preprint”[Publication Type]) AND (2020/1/1:2021/12/31[pdat])

## Acknowledgments

The authors thank Katherine Majewski for her invaluable expertise and support in coordinating the focus groups and survey, in addition to the creation of the NIH Preprint Pilot Toolkit and tutorial, Christopher Belter for his help with evaluation planning, and Jerry Sheehan for his feedback on this paper. We thank the NCBI Literature Program for implementing all the processes and presentation rules needed to support preprints in PMC and PubMed, including Kathi Canese, Jessica Chan, Chris Kelly, Erin Schmieder, Martin Latterner, Sergey Krasnov, Vladimir Sarkisov, and Andrei Kolotev. Thanks also to Ms. Schmieder for ongoing user engagement on preprints and comments on this paper and to Elizabeth Small for moderating focus group sessions. The authors are grateful to Yolanda L. Jones, NIH Library Editing Services, for editing assistance. Finally, special thanks to Becca Hohmann from PMC Operations for her management of the conversion and ingest processes for preprint records throughout Phase 1.

We also acknowledge the valuable service preprint servers provide and thank all those who provided email feedback, participated in focus groups, or responded to the survey.

