## Supplemental File 2 for "Phase 1 of the National Institutes of Health Preprint Pilot: Testing the viability of making preprints discoverable in PubMed Central and PubMed"

Appendix 2: NIH Preprint Pilot Survey Questions (2021)

OMB Control Number: **0925-0648**
Expiration Date: **06/30/2024**
Public reporting burden for this collection of information is estimated to average **10** minutes per response, including the time for reviewing instructions, searching existing data sources, gathering and maintaining the data needed, and completing and reviewing the collection of information. An agency may not conduct or sponsor, and a person is not required to respond to, a collection of information unless it displays a current valid OMB control number. Send comments regarding this burden estimate or any other aspect of this collection of information, including suggestions for reducing this burden, to NIH, Project Clearance Branch, 6705 Rockledge Drive, MSC 7974, Bethesda, MD 20892-7974, ATTN: PRA (0925-0648). Do not return the completed form to this address.

Q1 You were just viewing a **preprint** record in PubMed Central or PubMed. How clear was it to you that you were viewing a preprint?

- Unclear
- Somewhat clear
- Clear

Q2 Which category **best** defines your role when using PubMed or PubMed Central®?

- Student
- Researcher
- Librarian/Information Specialist
- Health Care Provider
- Journalist
- Educator
- Other

Q3 Had you heard of **preprints** before today?

- Yes
- No

Q4 What is your experience with preprints? (Select all that apply.)
 
I have:

- Read a preprint.
- Posted a preprint.
- Cited a preprint in the course of my work.
- Commented on a preprint in the course of my work.
- Shared a preprint with a colleague.
- Never used a preprint, but I know of them.
- Other: ________________________________________________

Q5 In a few words, how would you define a preprint?

________________________________________________________________

Q6 In your opinion, how important is it for the scientific community to be able to discover and access preprints?

- Extremely important
- Very important
- Moderately important
- Slightly important
- Not at all important

Q7 In your opinion, how important is it for the scientific community to be able to discover and access preprints about emerging topics like COVID-19?

- Extremely important
- Very important
- Moderately important
- Slightly important
- Not at all important

Q8 How important is it for **you** to be able to discover and access preprints for your work?

- Extremely important
- Very important
- Moderately important
- Slightly important
- Not at all important

Q9 How important is it for you to be able to discover and access preprints about emerging topics like COVID-19?

- Extremely important
- Very important
- Moderately important
- Slightly important
- Not at all important

Q10 How important is it for you to be able to discover and access preprints via PubMed or PubMed Central?

- Extremely important
- Very important
- Moderately important
- Slightly important
- Not at all important

Q11 The National Institutes of Health encourages the availability of preprints. Does this increase or decrease your trust in the information you find in **preprints**?

- Significantly decreases my trust
- Slightly decreases my trust
- Does not change my trust
- Slightly increases my trust
- Significantly increases my trust

Q12 To what extent does having preprints in PubMed or PubMed Central increase or decrease your trust in the information found in PubMed or PubMed Central?

- Significantly decreases my trust
- Slightly decreases my trust
- Does not change my trust
- Slightly increases my trust
- Significantly increases my trust

Q13 Please explain how having preprints **in PubMed or PubMed Central** affects your trust in **the information found in PubMed or PubMed Central**:

________________________________________________________________

Q14 What additional information about a preprint would help you understand or evaluate the information being reported?

________________________________________________________________

Q15  Do you have any other thoughts on preprints that would like to share?

________________________________________________________________

Q16 Which of the following activities do you do regularly?

- I read journal articles relevant to my interests.
- I search for journal articles related to specific research projects.
- I search for journal articles related to a specific disease or condition for personal use.

Q17 Which of the below activities have you done, even once?

- Peer-reviewed a paper or journal
- Authored a journal article

Q18 How confident are you in your ability to assess the scientific rigor and quality of a research article?

- Extremely confident
- Very confident
- Moderately confident
- Slightly confident
- Not at all confident

Q19 What indicators of scientific transparency do you use in evaluating scientific studies?

- Conflict of interest statement
- Availability of original data
- Funding statements
- Other ________________________________________________

End of Block: NIH Preprint Pilot Questionnaire
