## Supplemental File 1 for "Phase 1 of the National Institutes of Health Preprint Pilot: Testing the viability of making preprints discoverable in PubMed Central and PubMed"

### **Appendix 1: Focus Group Scripts (2021)**

***Standard Introduction:***

[Facilitator to give their standard introduction]

The reason we’re here today is to learn what people think about preprints. This is one of a few focus groups to hear from various target audiences. In brief, the National Library of Medicine started a project last June to test making preprints available via PubMed Central and cited in PubMed. The project is currently limited to preprints related to COVID. As it might be controversial to link preprints from a bibliographic database that is known for providing access to the published, peer-reviewed scientific literature, we’re very interested in what people think about preprints; how you use them; what questions you have, etc.

I’m going to lead our discussion today. I will be asking you questions and then encouraging and moderating our discussion.

This focus group will be recorded. The identities of all participants will remain confidential. The recording allows us to revisit our discussion for the purposes of developing research papers and presentations.

To allow our conversation to flow more freely, I’d like to go over some ground rules.

1. Only one person speaks at a time. It is difficult to hear everyone’s experience and perspective on our web meeting if there are multiple voices at once.
2. Everyone doesn’t have to answer every single question, but I’d like to hear from each of you today as the discussion progresses.
3. This is a confidential discussion. Names of participants will not be included in the final report. It also means, except for the report that will be written, what is said in this session stays in this session.
4. We stress confidentiality because we want an open discussion. We want all of you to feel free to comment on each other’s remarks without fear your comments will be repeated later and possibly taken out of context.
5. There are no “wrong answers,” just different opinions. Say what is true for you, even if you’re the only one who feels that way. Don’t let the group sway you. But if you do change your mind, let me know.
6. Let me know if you need a break.
7. Are there any questions?

Before we start, I’d like to know a little about each of you. Please tell us your name, how long you’ve been a clinician, and what your area of clinical expertise.

***Tailored Questions***

Each focus group had questions specifically tailored to their role.

##### Research Librarians

*Value and Assessment*

1. How do you assess a research article?
   a. What characteristics of the research do you look for?
   b. What characteristics of the publication	do you look for?
2. Have your criteria or process for evaluating research publications changed over time? If so, how?
3. Do you think preprints are valuable? Why or why not?
4. How do you assess a preprint?
   a. What characteristics of the publication do you look for?
   b. Is your process for assessing preprints different from your process for assessing other forms of research reporting? If so, in what ways?
5. Has COVID changed your perception of the value of preprints? If so, how?
6. Do you look for response or comments from the author’s peers on a preprint?
   a. Where do you look?
   b. Do comments affect the value of a preprint or the research? If so, in what ways?

*Search and Retrieval*

1. How do you find preprints?
2. Do you seek preprints out separately? If so, in what circumstances?
   a. If so, where do you look?
   b. Should seeking preprints be a separate step?
3. Has COVID changed whether and how you seek out preprints? In what ways?
4. How often do you follow up use of a preprint by seeking out the published version? Why?

*Use*

1. Have you ever used the information from a preprint in decision-making? If so, what was your experience?
2. What factors influence you (or would influence you) to use information found in a preprint?

*Dissemination*

1. Have you ever shared a preprint or information from a preprint with others? If so, what was your experience?
2. What factors influence you (or would influence you) in deciding to share a preprint or information found in a preprint?
3. In what circumstances would you publicly cite a preprint as a source? Why?
4. How might you report the contents of a preprint differently from a published article?

*Education*

1. How and from whom have you learned about preprints?
2. Has what you learned changed your perception of preprints or research reporting?
3. Has what you learned changed whether and how you search or evaluate the literature? In what ways?
4. Have you ever taught others about preprints? If so, what did you learn in the process?
5. What should researchers to know about preprints?
6. What should clinicians know about preprints?
7. What should librarians know about preprints?
8. What should journalists know about preprints?

*NIH/NLM Roles and Responsibilities*

1. What is NLM’s responsibility in collecting, describing, and providing access to preprints?
2. Is it appropriate that NIH allows preprints to be cited in grant applications and progress reports? Why or why not?
3. Does the NIH Preprint Pilot increase or decrease your trust in NIH? In NLM? In what ways?
4. How does the availability of preprints in PubMed Central affect how you value preprints?
   a. How does the availability of preprints in other places (e.g., SCOPUS) affect how you value preprints?
5. What is the most important information about a preprint (e.g., metadata) that NLM should be providing, along with the preprint?
6. What actions can NLM or other libraries take to improve research quality and research reporting?
7. Is there anything else you would like NLM to know about preprints?

##### Biomedical Researchers

*Value and Assessment*

7. How do you assess a research article?
   a. What characteristics of the research do you look for?
   b. What characteristics of the publication	do you look for?
8. Have your criteria or process for evaluating research publications changed over time? If so, how?
9. Do you think preprints are valuable? Why or why not?
10. How do you assess a preprint?
    a. What characteristics of the publication do you look for?
    b. Is your process for assessing preprints different from your process for assessing other forms of research reporting? If so, in what ways?
11. Has COVID changed your perception of the value of preprints? If so, how?
12. Do you look for response or comments from the author’s peers on a preprint?
    a. Where do you look?
    b. Do comments affect the value of a preprint or the research? If so, in what ways?

*Search and Retrieval*

1. How do you find preprints?
2. Do you seek preprints out separately? If so, in what circumstances?
   a. If so, where do you look?
   b. Should seeking preprints be a separate step?
3. Has COVID changed whether and how you seek out preprints? In what ways?
4. How often do you follow up use of a preprint by seeking out the published version? Why?

*Use*

1. Have you ever used the information from a preprint in decision-making? If so, what was your experience?
2. What factors influence you (or would influence you) to use information found in a preprint?

*Dissemination*

1. Have you ever shared a preprint or information from a preprint with others? If so, what was your experience?
2. What factors influence you (or would influence you) in deciding to share a preprint or information found in a preprint?
3. In what circumstances would you publicly cite a preprint as a source? Why?
4. How might you report the contents of a preprint differently from a published article?

*Education*

1. How and from whom have you learned about preprints?
2. Has what you learned changed your perception of preprints or research reporting?
3. Has what you learned changed whether and how you search or evaluate the literature? In what ways?
4. Have you ever taught others about preprints? If so, what did you learn in the process?
5. What should researchers to know about preprints?
6. What should librarians know about preprints?

*Production*

1. Have you ever posted a preprint? If so, what was your experience?
   a. Was your intention to publish, following the posting of the preprint? Why or why not?
2. Why might you post a preprint? What are the benefits?
3. Do you make your original data and data analysis available to the public? Why or why not?
4. Are you likely to change the content of a report based on peer feedback from your preprint?
   a. What kinds of feedback are most likely to influence you to change how you describe your research results?
5. What concerns might you have about posting a preprint? What are the risks?
6. Have you commented on someone else’s preprint? What was your experience? What did you expect to happen, based on your comments?

*NIH/NLM Roles and Responsibilities*

1. What is NLM’s responsibility in collecting, describing, and providing access to preprints?
2. Is it appropriate that NIH allows preprints to be cited in grant applications and progress reports? Why or why not?
3. Does the NIH Preprint Pilot increase or decrease your trust in NIH? In NLM? In what ways?
4. How does the availability of preprints in PubMed Central affect how you value preprints?
   a. How does the availability of preprints in other places (e.g., SCOPUS) affect how you value preprints?
5. What is the most important information about a preprint (e.g., metadata) that NLM should be providing, along with the preprint?
6. What actions can NLM or other libraries take to improve research quality and research reporting?
7. Is there anything else you would like NLM to know about preprints?

Healthcare Journalists

*Value and Assessment*

1. How do you assess a research article?
   a. What characteristics of the research do you look for?
   b. What characteristics of the publication	do you look for?
2. Have your criteria or process for evaluating research publications changed over time? If so, how?
3. Do you think preprints are valuable? Why or why not?
4. How do you assess a preprint?
   a. What characteristics of the publication do you look for?
   b. Is your process for assessing preprints different from your process for assessing other forms of research reporting? If so, in what ways?
5. Has COVID changed your perception of the value of preprints? If so, how?
6. Do you look for response or comments from the author’s peers on a preprint?
   a. Where do you look?
   b. Do comments affect the value of a preprint or the research? If so, in what ways?

*Search and Retrieval*

1. How do you find preprints?
2. Do you seek preprints out separately? If so, in what circumstances?
   a. If so, where do you look?
   b. Should seeking preprints be a separate step?
3. Has COVID changed whether and how you seek out preprints? In what ways?
4. How often do you follow up use of a preprint by seeking out the published version? Why?

*Use*

1. Have you ever used the information from a preprint in decision-making? If so, what was your experience?
2. What factors influence you (or would influence you) to use information found in a preprint?

*Dissemination*

1. Have you ever shared a preprint or information from a preprint with others? If so, what was your experience?
2. What factors influence you (or would influence you) in deciding to share a preprint or information found in a preprint?
3. In what circumstances would you publicly cite a preprint as a source? Why?
4. How might you report the contents of a preprint differently from a published article?

*Education*

1. How and from whom have you learned about preprints?
2. Has what you learned changed your perception of preprints or research reporting?
3. Has what you learned changed whether and how you search or evaluate the literature? In what ways?
4. What should journalists know about preprints?

*NIH/NLM Roles and Responsibilities*

1. What is NLM’s responsibility in collecting, describing, and providing access to preprints?
2. Is it appropriate that NIH allows preprints to be cited in grant applications and progress reports? Why or why not?
3. Does the NIH Preprint Pilot increase or decrease your trust in NIH? In NLM? In what ways?
4. How does the availability of preprints in PubMed Central affect how you value preprints?
   a. How does the availability of preprints in other places (e.g., SCOPUS) affect how you value preprints?
5. What is the most important information about a preprint (e.g., metadata) that NLM should be providing, along with the preprint?
6. What actions can NLM or other libraries take to improve research quality and research reporting?
7. Is there anything else you would like NLM to know about preprints?

##### Clinicians

*Value and Assessment*

1. How do you assess a research article?
   a. What characteristics of the research do you look for?
   b. What characteristics of the publication	do you look for?
2. Have your criteria or process for evaluating research publications changed over time? If so, how?
3. Do you think preprints are valuable? Why or why not?
4. How do you assess a preprint?
   a. What characteristics of the publication do you look for?
   b. Is your process for assessing preprints different from your process for assessing other forms of research reporting? If so, in what ways?
5. Has COVID changed your perception of the value of preprints? If so, how?
6. Do you look for response or comments from the author’s peers on a preprint?
   a. Where do you look?
   b. Do comments affect the value of a preprint or the research? If so, in what ways?

*Search and Retrieval*

1. How do you find preprints?
2. Do you seek preprints out separately? If so, in what circumstances?
   a. If so, where do you look?
   b. Should seeking preprints be a separate step?
3. Has COVID changed whether and how you seek out preprints? In what ways?
4. How often do you follow up use of a preprint by seeking out the published version? Why?

*Use*

1. Have you ever used the information from a preprint in decision-making? If so, what was your experience?
2. What factors influence you (or would influence you) to use information found in a preprint?

*Dissemination*

1. Have you ever shared a preprint or information from a preprint with others? If so, what was your experience?
2. What factors influence you (or would influence you) in deciding to share a preprint or information found in a preprint?
3. In what circumstances would you publicly cite a preprint as a source? Why?
4. How might you report the contents of a preprint differently from a published article?

*Education*

1. How and from whom have you learned about preprints?
2. Has what you learned changed your perception of preprints or research reporting?
3. Has what you learned changed whether and how you search or evaluate the literature? In what ways?
4. What should clinicians know about preprints?

*NIH/NLM Roles and Responsibilities*

1. What is NLM’s responsibility in collecting, describing, and providing access to preprints?
2. Is it appropriate that NIH allows preprints to be cited in grant applications and progress reports? Why or why not?
3. Does the NIH Preprint Pilot increase or decrease your trust in NIH? In NLM? In what ways?
4. How does the availability of preprints in PubMed Central affect how you value preprints?
   a. How does the availability of preprints in other places (e.g., SCOPUS) affect how you value preprints?
5. What is the most important information about a preprint (e.g., metadata) that NLM should be providing, along with the preprint?
6. What actions can NLM or other libraries take to improve research quality and research reporting?
7. Is there anything else you would like NLM to know about preprints?
